# Effects of stand characteristics on tree species richness in and around a conservation area of northeast Bangladesh

**DOI:** 10.1101/044008

**Authors:** Muha Abdullah Al Pavel, Sharif A. Mukul, Mohammad Belal Uddin, Kazuhiro Harada, Mohammed A. S. Arfin Khan

## Abstract

We investigated the effect of tree cover, forest patch and disturbances on tree species richness in a highly diverse conservation area of northeast Bangladesh. A systematic sampling protocol was adopted and 80 sub-plots from twenty five 1 ha plots were used for the vegetation survey. Linear regression analysis was performed to understand the effect of patch area, disturbances and tree cover on tree species richness. Ordination using Redundancy analysis (RDA) and Non-metric Multi Dimensional Scaling (NMDS) were also performed to explore the tree species compositional similarities along the stand characteristics gradient and locations of the sample plots. Our study revealed that, forest patch size has greater influence on species richness. Areas with medium level of disturbances have shown greater species richness. In constrained ordination the selected explanatory variables regulated the richness of common species. Our findings can be useful for better forest management and restoration of landscapes of conservation needs using ecologically important species.

## 1 Introduction

Biodiversity has become the issue of global attention because of growing awareness of its importance on the one hand and its rapid depletion worldwide (Shrestha 1999; Singh 2002). Apart from its immense economic and aesthetic value, biodiversity is essential for proper functioning of ecosystems and its stability (Ehrlich and Wilson 1991; Holdgate 1996; Tilman 2000; Singh 2002). In the tropics, biodiversity is threatened by land use change, anthropogenic disturbances and more recently by warming climate (Singh 2002; Alamgir et al. 2015). Disturbances created by factors like fire, habitat loss, over exploitation, shifting cultivation and non-native species introduction influence forest dynamics and determine the tree species richness (Burslem and Whitmore 1999; Uddin et al. 2013; Mukul and Herbohn 2016).

Bangladesh is one of the worlds’ most densely populated countries experiencing intensive pressure on forest areas through land use change and human disturbance (Mukul et al. 2012; Uddin et al. 2013; Mukul 2014; Sohel et al. 2015). The country has a forest over of about 14.4 million hectares representing 17.1% of the total land area (Khan et al. 2007; GOB 2010). Moreover, 90% of the country’s forests are degraded with an annual rate of forest loss of about 0.3% in the country (Chowdhury and Koike, Kibria et al. 2011; Rahman et al. 2015). One of the major causes of biodiversity loss in the country is fragmentation of large blocks of natural forests into small and sometimes isolated patches of native vegetation surrounded by a matrix of agricultural and/or other land-use (see Kibria et al. 2011; Rahman et al. 2013; Uddin et al. 2013; Mukul 2014). Forest patches, therefore, provide an abrupt edge between forests and surrounding vegetation in the country (Murcia 1995). These edges have serious ecological and environmental consequences, such as elimination of native species and development of degraded landscape dominated by invasive species from surrounding areas (Pandey et al. 2003; Uddin et al. 2013). In fact, processes involved in tree regeneration can be influenced by many factors, like variations in the seed dispersal intervals, seed quality, wind direction, wind speed, slope of the site, gradients, aspects, soil moisture availability etc. (Vieira and Scariot 2006). Moreover, fire, light intensity and tree-fall rates are higher on edges of recent fragments which promotes invasion by weeds, pioneer species and vines (Laurance and Bierregaard 1997; Brokaw 1998).

Although, disturbances both natural and human made are common in the country, and cotinually shaping plant species richness, composition and diversity in the country, there are only a few studies that systematically investigate this aspect in the country’s forest (Uddin et al. 2013; Mukul 2014). Against this backdrop, the present study investigated the effects of stand characteristics on plant species richness in one of the country’s most diverse forest patch – Lawachara National Park (LNP). The specific objective of the study was to assess the effect of patch size, disturbances and tree cover on tree species richness at the landscape level in LNP. The finding of this study could be useful to make more efficient management plans to support the conservation of native biodiversity in country’s remaining forests. Moreover, the study could be helpful to design future research on biodiversity pattern at different landscapes and disturbance gradient in other forest areas of Bangladesh.

## 2 Materials and Methods

### 2.1 The study site

LNP lies between 24°30′ – 24°32′ N and 91°37′ – 91°47′ E, and is one of the oldest protected areas of the country (Figure 1; Mukul et al. 2014). Administratively, it is under the Maulivibazar forest division and located nearly 160 km northeast of Dhaka, the capital city of Bangladesh. The park covers an area of about 1250 ha (12.5 km^2^). The climate is generally warm and humid, but cool and pleasant during the winter. The temperature varies with an average from a minimum of 27°C in February to maximum of 36°C during June. The humidity is high throughout the year, with monthly average humidity varying from 74% in March to 89% in July. The park is situated in the wettest region in the country with an annual average rainfall of 4000 mm per year (Mukul 2008). Geographically, the area is undulating with gentle slopes and elevation ranging between 0 – 50 m (Mukul et al. 2014). The soil of the forest can be categorized as hill brown sandy loams with slight to strong acidity (NSP 2006). There are shallow over sandstone bedrocks on high hills and accumulation of humus in the top soil is small due to rapid decomposition of debris under moist warm tropical conditions (NSP 2006).

**Figure 1.**
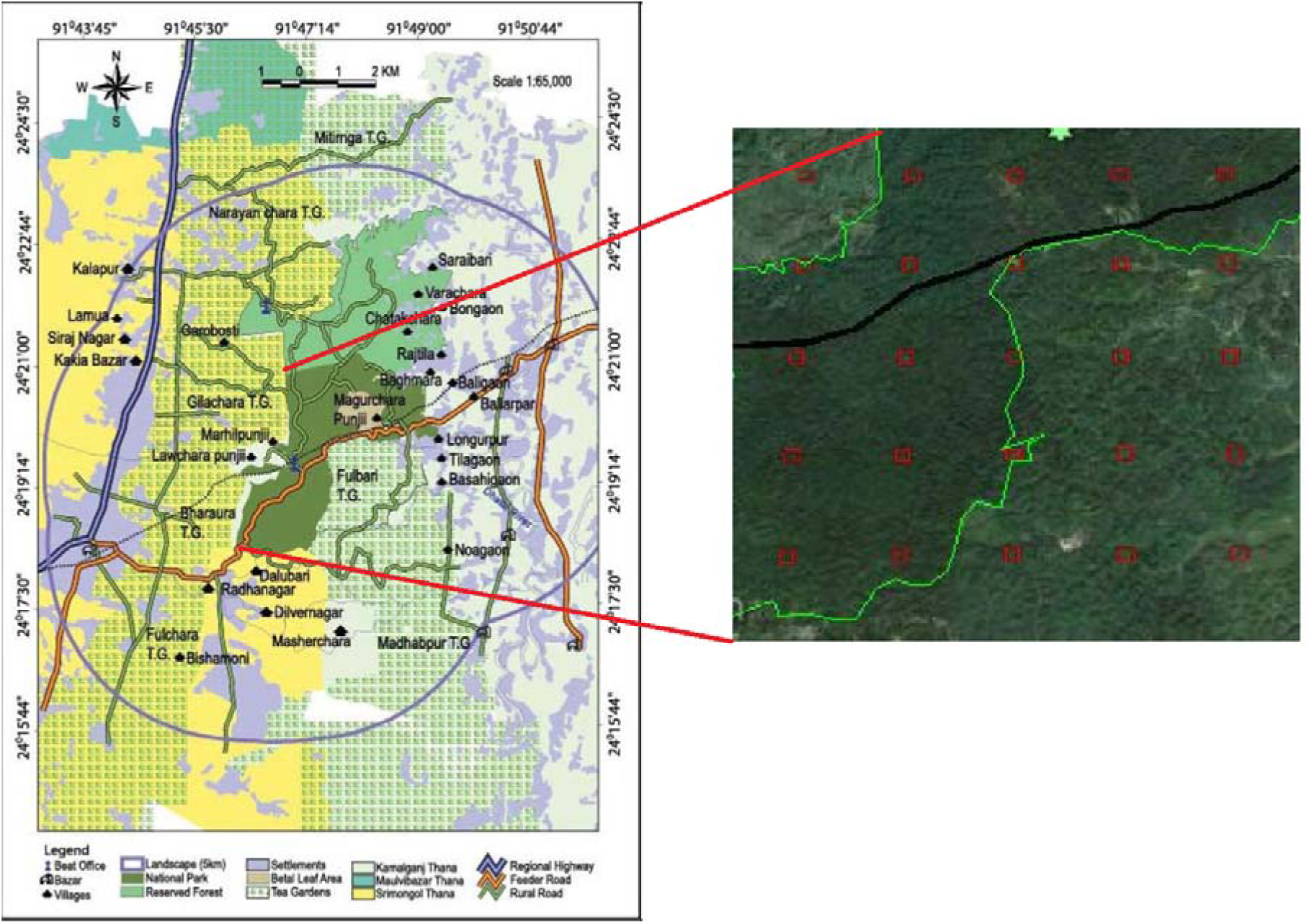
Land use/cover map of Lawachara National Park and adjoining area (left), and the location of the survey plots (right)

### 2.2 Sampling protocol

A systematic sampling design was adopted for the study (Bongers et al. 2009). Twenty five square plots of 100 m × 100 m (1 ha) size were used for the vegetation survey. Sample plots were established within a 4 km × 4 km grid area with 800 m spacing from one plot to another. Each square plot was divided into sub-plots with unique forest type (forest patch). Forest patches were selected with respect to their type, form, shape, forest area percentage and number of vegetation corridors within a forest patch (Galanes and Thomlinson 2008). Forest patches used in this study were homogenous in nature and differs from the surrounding vegetation (Forman 1995). A Google map of the area was created to select and lay out the sample plots. The location of the first plot was chosen randomly, and the locations of the remaining plots were fixed regularly based on the position of the first plot as described in Acharya (1999). Location of the plots within LNP was then traced using a hand-held GPS (Model: Garmin, USA). The sample plots cover about 2% of the total area of LNP. Within each plot all mature tree ≥ 5 cm were identified and measured at the diameter at breast height (dbh; 1.37 meter). All tree seedlings having a diameter at collar region < 2.5 cm and a height < 1 m, and all tree saplings with dbh greater than 2.5 cm but less than 5 cm were also recorded. In case of disturbances, the number of visible disturbances present or absent was recorded by visual interpretation and through a checklist questionnaire. Also, patch area and tree cover were calculated as the percentage in sample plot cross with map respectively through direct field observation that was the effect of species diversity (Rao et al. 1990; Raghubanshi and Tripathi 2009).

### 2.3 Data analysis

We used species richness and Shannon-Wiener index to describe the species diversity (Whittaker 1972). Species richness (*N*) was measured as the absolute number of unique species in each sample plot, Shanon-Wienner index (*H′*) was calculated as: *H*′ = -Σ*Pi* In *Pi*; where, *Pi* is the relative importance value of species. We, however, only used species richness for detail statistical analysis. We performed linear regression analysis to find out the effect of patch area, tree cover and disturbances on species richness. Analysis of variance (ANOVA) was also performed to test the significance of each linear model. A checklist was developed after Uddin et al. (2013) to identify forest disturbances and their relative influence in the study plots. These parameters were given a value ranging between 0 to 8, with 8 being the highest and 0 being the lowest (Biohabitats 2012). The main advantage of using score methods in evaluating representativeness was that they summarized information about the range in variation. Interaction of disturbance attributes or score of total disturbance: *F*\**S*\**D*\**I*, where ‘*F*’ is the Frequency, ‘*S*’ is Size, ‘*D*’ is the Duration and ‘*I*’ is the Intensity (Supplementary material 1). We used statistical package Biodiversity–R (Vegan 1.17-4 package) and R (version 2.10.1) for our multivariate analysis. Both constrained (RDA) and unconstrained (NMDS) ordination were performed to see species compositional similarities where in constrained ordination species composition was analysed along stand characteristics gradient and in unconstrained ordination along the position of the plots within the study area in LNP.

## 3 Results

### 3.1 Species richness and diversity

Altogether we recorded 848 individuals representing 130 tree species from our 80 sub-plots (Annex 1). *Artocarpus chaplasha* was the most common species (35 individuals) in the study area. Other common species were — *Aphanamixis polystachya, Albizia lebbeck, Taphrosia candida, Ilex godajam* etc. Species richness and Shannon-Weiner index in sub-plots of different location within the study area are listed in Table 1.

**Table 1.** Species richness and Shannon-Weiner index in different study plots in the area

| Parameter | Core area* | Forest edge | Surrounded area |
| --- | --- | --- | --- |
| Number of sub-plot | 32 | 12 | 36 |
| Species richness ($N$) | 15.96(±9.71) | 12.25(±7.53) | 10.00(±7.52) |
| Shanon-Weinner's Index ($H'$) | 2.47(±0.93) | 2.18 (±1.01) | 1.91(±1.04) |

\*values in the parenthesis represent standard deviation.

### 3.2 Effect of patch size on tree species richness

Linear regression using species richness and patch area showed a significant (*p*<0.05) positive relationship. For trees, the value of R–squared (30%) means that the model only explains a fraction of the total variance in species richness data. The adjusted R-squared value (29%) was as close as R-squared value that provided additional information of the variance in LNP (Table 2). In addition, for sapling, the value of R-squared (35%) means that the model only explained a fraction of the total variance in species richness data which was less than trees regression model. In contrast, the adjusted R-squared value (34%) was as close as R-squared value which provided additional information of the variance in LNP (Table 2). In case of seedling, the value of R-squared (35%) means that the model only explained a fraction of the total variance in species richness data. Alternatively, the adjusted R-squared value (35%) was as close as R-squared value which provided additional information of the variance in LNP (Table 2). Furthermore, result of ANOVA showed of the quality effect of patch area on species richness in LNP. For the tree species richness, the *F* statistic value (*F*=34.39) provided slight evidence that the model explained a few of the variance. It was also noticed that the significance level (*p*<0.05) was strong, but slightly evident that the patch area explained the variation in species richness in LNP. Once again that this significance level was the same as calculated by earlier tests (*p*<0.05), indicating strong evidence that effect of patch area contributes to explaining of species richness in LNP (Annex 2; Table A). In case of sapling species richness, on the other hand, the *F* statistic value (*F*=43.42) afforded moderate evidence that the model explained some of the variance. It noticed that the significance level (*p*<0.05) was indicated strong, but moderate evidence that effect of patch area contributes to explaining on species richness of variance in LNP (Annex 2; Table A). Furthermore, for the seedling species richness, the *F* statistic value (F=43.70) has sharp evidence that the model explained some of the variance than previous indicating *F* value. And, it noticed that the significance level (*p*<0.05) was indicated strong, but sharply evident that effect of patch area contributes to explaining on the seedling species richness of variance in LNP (Annex 2; Table A). Similarly, since the linear regression model fits a straight line that attempted to get as close as possible to the observed values, compared the predictions with the actual observations. And, this fits a straight line showed (35.9%; R-squared) of the total variance for seedling species followed by sapling species (35.8%; R-squared) and tree species (30.6%; R-squared). Besides, the dashed lines explained where the regression line was at 95% confidence (Figure 2).

**Table 2.** Linear regression analysis of species richness vs. patch area, disturbance and tree cover

| Explanatory variable | Dependent variable | Intercept (a) | Co-efficient (b) | Adjusted $R^2$ | SE |
| --- | --- | --- | --- | --- | --- |
| Patch area | Trees | 6.48 | 0.18 | 0.29 | 7.39 |
|  | Sapling | 2.38 | 0.14 | 0.34 | 5.21 |
|  | Seedling | 4.74 | 0.16 | 0.35 | 5.13 |
| Disturbance | Tree | -9.20 | 0.16 | 0.67 | 5.03 |
|  | Sapling | -3.91 | 0.08 | 0.32 | 5.21 |
|  | Seedling | -0.77 | 0.08 | 0.24 | 6.28 |
| Tree cover | Tree | 8.23 | 0.12 | 0.14 | 8.14 |
|  | Sapling | 3.39 | 0.10 | 0.21 | 5.64 |
|  | Seedling | 5.88 | 0.11 | 0.21 | 6.41 |

SE= Std. error of the estimate

**Figure 2.**
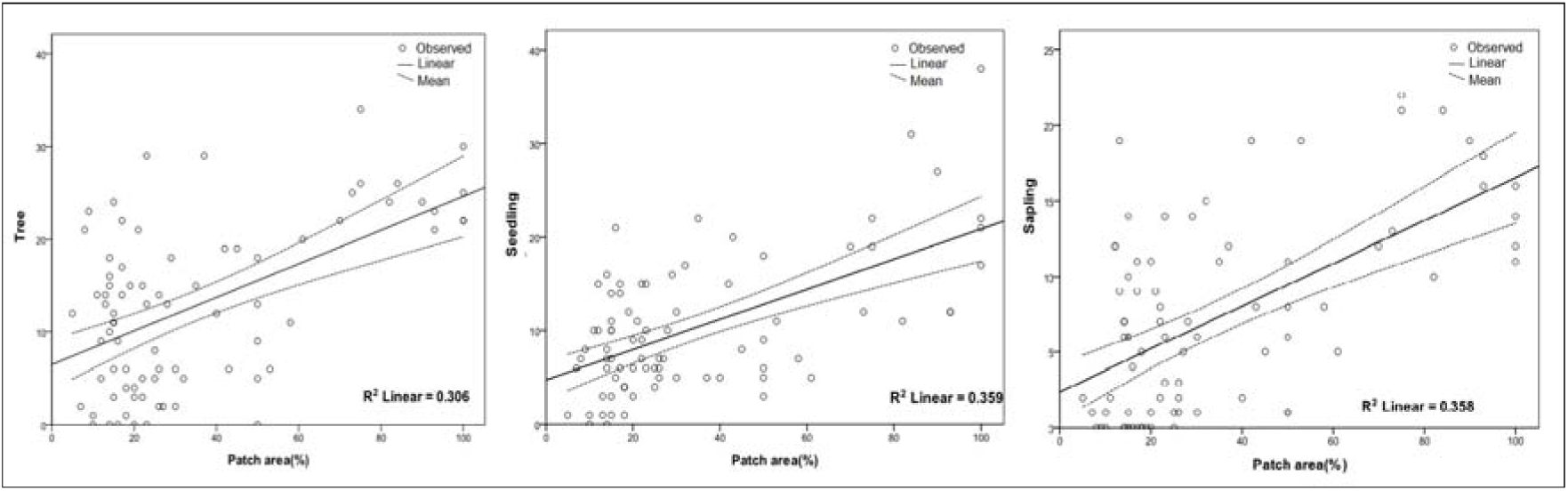
Effect of patch area (%) on species richness. Here is observed values (circles) and predicted values (connected by solid line) for the linear regression. The dotted lines show the 95% confidence interval for the mean of observations.

### 3.3 Effect of disturbance on species richness

Linear regression model of disturbance showed a significant positive relationship with species richness. For tree, the value of R-squared (67%) means that the model only explained moderately a fraction of the total variance in species richness data. And then, the adjusted R-squared value (67%) was as close as R-squared value which provided extra information of the variancein LNP (Table 2). In case of sapling, however, indicated positive, but a sharply linear relationship with disturbance parameter. The value of R-squared (33%) means that, the model explained sharply a fraction of the total variance in species richness data which was more than trees regression model R-squared value. In contrast, the adjusted R-squared value (32%) was as close as R-squared value (33%) which provides little additional information of the variance which was also more than trees regression model R-squared value in LNP (Annex 2; Table B). In case of the seedlings, there was a positive, but slightly linear relationship with disturbance pattern. The value of R-squared (25%) means that the model only explained slight a fraction of the total variance in species richness data. Alternatively, the adjusted R-squared value (24%) was as close as R-squared values (25%) which afford extra information of the variance in LNP (Table 2). Similarly, the outcome of ANOVA showed an expression of the quality effect of disturbance pattern on species richness in LNP. For the trees, the *F* statistic value (*F*=164.29) was sharply evidence that the model explained more of the variance. It was also noticed that the significance level (*p*<0.05) indicated strong evidence that effect of disturbance parameter contributes to explaining the variation in species richness in LNP. Once again that this significance level was the same as the significance level calculated by earlier tests (*p*<0.05) indicating strong evidence that effect of disturbance contributes to explaining of species richness in LNP (; Table 2). In case of saplings, the *F* statistic (*F*=39.47) was moderately evident that the model explained some of the variance. Similar to the trees, the significance level (*p*<0.05) indicated strong evidence that the effect of disturbance contributes in explaining the variation in species richness (Table 2). For the seedling species richness, the *F* statistic (*F*=26.91) was slightly evident that the model explained some of the variance than previous indicating *F* value. And then, it was noticed that the significance level (*p*<0.05) indicated strong, but slight evidence that effect of disturbance contributes to explaining on the seedling species richness of variance data (Annex 2; Table B). Likewise, since the linear regression model fits a straight line that attempted to get as close as possible to the observed values, compared the predictions with the actual observations. This fits a straight line showed 67.8% (R-squared) of the total variance for tree species followed by sapling species (33.8%; R-squared) and seedling species (25.7%; R-squared). The dashed lines illustrated where the regression line was at 95% confidence (Figure 3).

**Figure 3.**
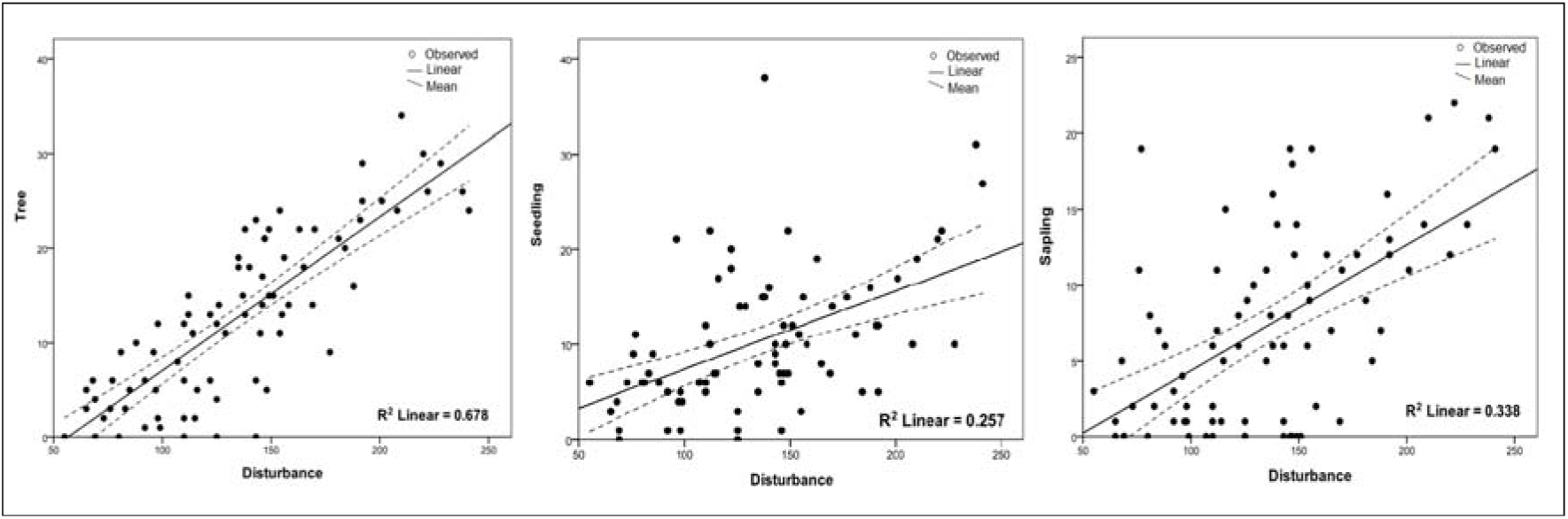
Effect of disturbance on species richness. Here is observed values (circles) and predicted values (connected by solid line) for the linear regression. The dotted lines show the 95% confidence interval for the mean of observations.

### 3.4 Effect of tree covers on species richness

Result of linear regression of tree cover showed a significant positive relationship with species richness in LNP. For trees, the value of R-squared (15%) means that the model only explained a fraction of the total variance in species richness. The adjusted R-squared value (14%) was as close as R-squared value which provided little additional information about the variance (Table 2). For sapling, indicated positive, but a moderate linear relationship with tree cover. The value of R-squared (22%) means that the model only explained a fraction of the total variance in species richness data which were more than trees regression model. In contrast, the adjusted R-squared value (21%) was as close as R-squared value that result provides extra information of the variance which was also more than trees regression model in LNP (Annex 2; Table C). In case of seedling, on the contrary, indicated positive, but a slightly linear relationship with tree cover. The value of R-squared (22%) means that the model only explained a fraction of the total variance in species richness data. Otherwise, the adjusted R-squared value (21%) was as close as R-squared value which provided extra information of the variance in LNP (Annex 2; Table C). Moreover, result of ANOVA showed an expression of the quality effect of tree cover on species richness in LNP. For the trees, the *F* statistic value (*F*=14.56) which was slightly evident that the model explained a few of the variance. It could be noticed also that the significance level (*p*<0.05) was indicated strong, but slightly evident that effect of tree cover contributes to explaining on species richness of variance data in LNP (Annex 2; Table C). Once again that this significance level was the same as the significance level calculated by earlier tests (*p*<0.05), indicating strong, evidence that effect of tree cover contributes to explaining of species richness in LNP (Table 2). In case of sapling, on the other hand, the *F* statistic value (*F*=22.42) which was slightly evidence that the model explained some of the variance. It could be noticed that the significance level (*p*<0.05) was indicated strong, but moderate evidence that effect of tree cover contributes to explaining on species richness of variance data in LNP (Table 2). For the seedling, the *F* statistic value (*F*=22.70) which was moderately evidence that the model explained some of the variance than previous indicating *F* value. And, it could be noticed that the significance level (*p*<0.05) was indicated strong, but sharply evidence that effect of tree cover contributes to explaining on the seedling species richness of variance data (Table 2). Equally, since the linear regression model fits a straight line that attempted to get as close as possible to the observed values, compared the predictions with the actual observations. And, this fits a straight line showed 22.5% (R-squared) of total variance for sapling species followed tree by species (22.3%; R-squared) and seedling species (25.7%; R-squared). Moreover, the dashed lines illustrated where the regression line was at 95% confidence (Figure 4).

**Figure 4.**
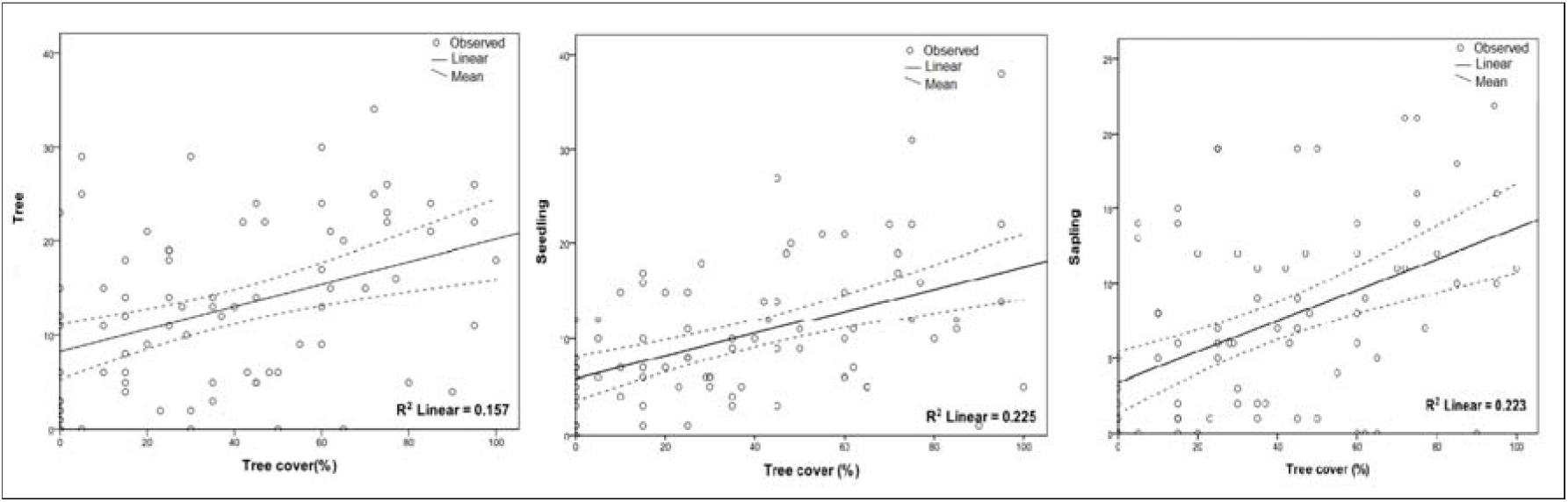
Effect of tree cover (%) on species richness. Here is observed values (circles) and predicted values (connected by solid line) for the linear regression. The dotted lines show the 95% confidence interval for the mean of observations.

### 3.5 Species compositional similarities

In RDA ordination total constrained and unconstrained variance was found 13.68% and 86.32% from total variance of 100%. RDA axis 1, 2 and 3 individually explained 80%, 10% and 8% of total constrained variance (Table 3). The percentage of variance of axis 1 and axis 2 were higher than the rest of the axes. In addition, permutation tests for all constrained eigenvalues based on 10000 permutations (under reduced model) revealed that RDA ordination was statistically significant (Pseudo *F*: 4.01), *p*<0.001. In contrast, RDA advance analyzed the results of PCA. It was constrained by disturbance variables and reflected ecological disturbance (Bray-Curtis distance). RDA ordination graph using scaling 1 (inter-sample distance) revealed that most of the sites were similar but few sites were showing high variability due to the influence of disturbance, patch area and tree cover percentage variable. Also, it showed that variables strongly regulated the richness of most certain species whereas, variables were not influenced for the distribution of few species used in this research. Moreover, the first axis described the disproportionately large number of species richness irrespective of the disturbance, patch area percentage and tree cover percentage variable (Figure 5). Similarly, when scaling 2 (inter-species) was used with environmental matrix, RDA ordination graph showed that disturbance, patch area percentage and tree cover percentage variables regulated the richness of most certain species were not influences distribution of few species in the study area. Moreover, the first axis described the proportionately large number of patch site with richness irrespective of the disturbance; patch area percentage and the tree cover percentage variables (Figure 6). Moreover, disturbance level, patch area percentage and tree cover percentage were negatively correlated with RDA1 and RDA2 was also negatively correlated with disturbance and the tree cover percentage, but the patch area percentage was positive (Table 3).

**Table 3.** Variables and RDA ordination results

| Variable | RDA Axes |  |  |
| --- | --- | --- | --- |
|  | Axes 1 | Axes 2 | Axes 3 |
| Disturbance (%) | -0.92 | -0.22 | 0.30 |
| Patch area (%) | -0.73 | 0.59 | -0.31 |
| Tree cover (%) | -0.54 | -0.23 | -0.80 |
| Eigen value | 1.16 | 0.15 | 0.12 |
| Proportion explained | 0.80 | 0.10 | 0.08 |

**Figure 5.**
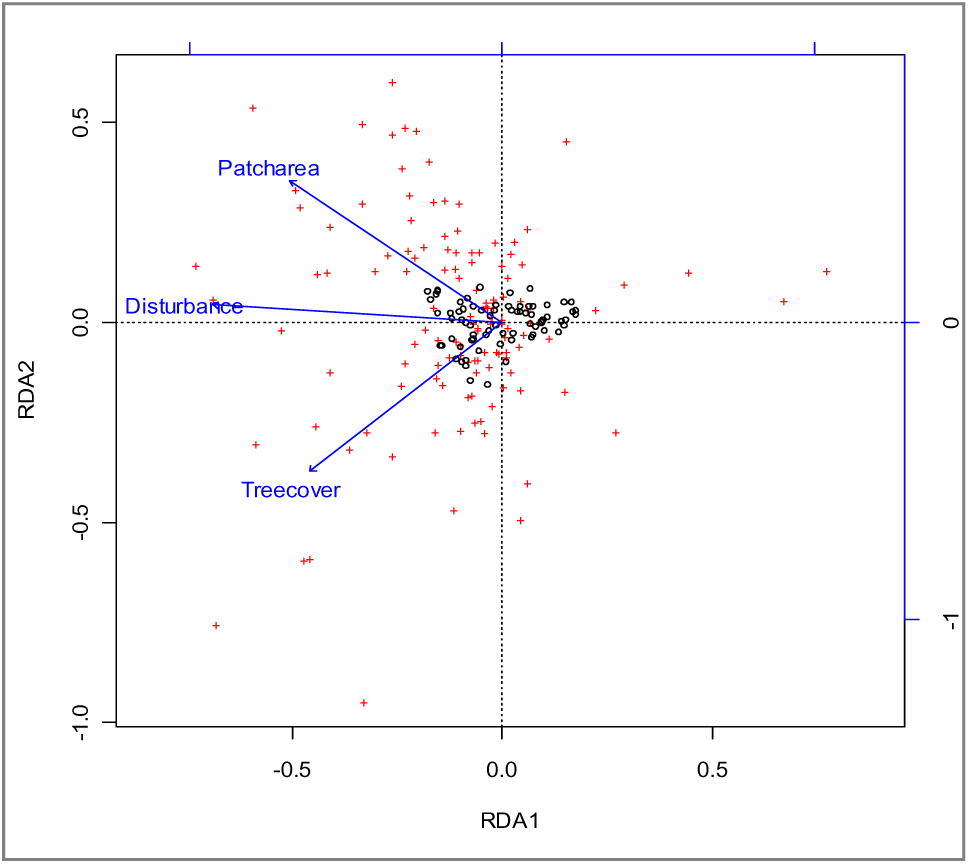
Species compositional similarities along the stand characteristics gradient and sites of differen location in the study area using redundancy analysis (RDA) scaling method one. Here is species (+) and plot (o).

**Figure 6.**
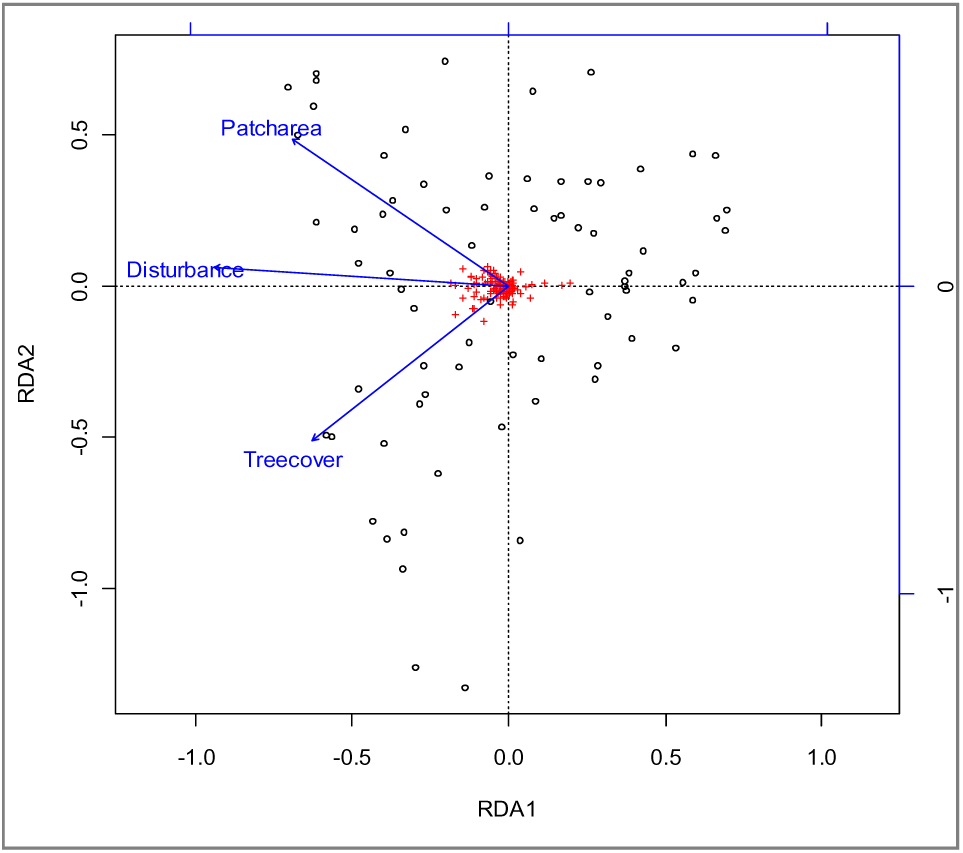
Species compositional similarities along the stand characteristics gradient and sites of differen location in the study area using redundancy analysis (RDA) scaling method two. Here is species (+) and plot (o).

Ordination using non-metric multi-dimensional scaling and using the Bray-Curtis distance and 1000 permutations showed a closer relationship in species composition between plots located in the core conservation area, followed by in forest edge and in the surrounding areas (Figure 7).

**Figure 7.**
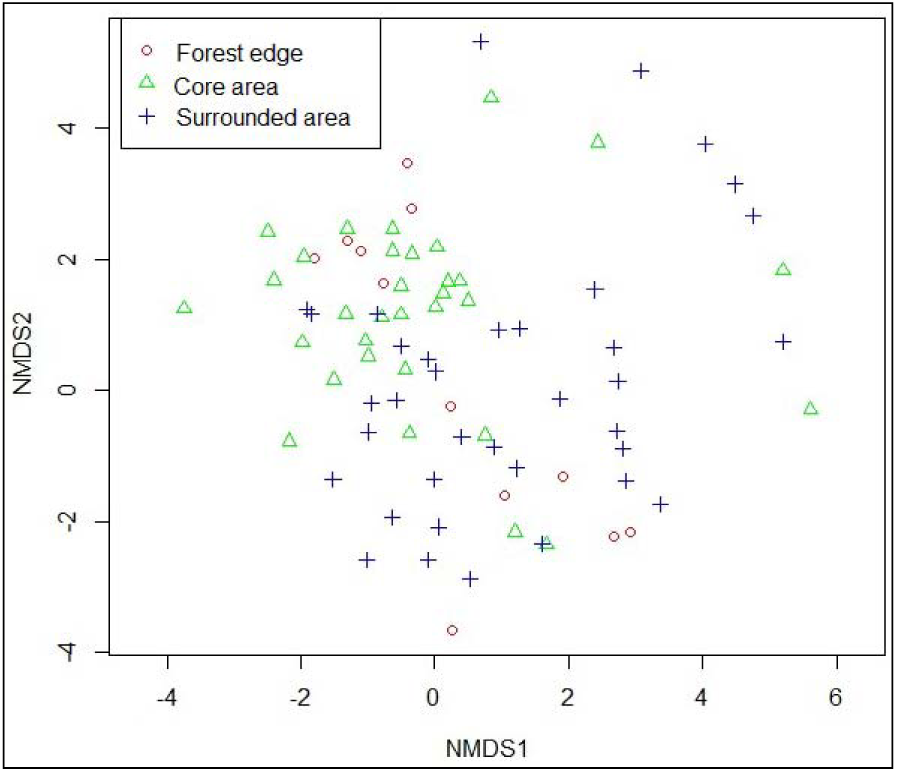
Species compositional similarities among sites of differen location in the study area using non-metric multi dimensional scaling and Bray-Curtis distance.

## 4 Discussion

We found stand characteristics as an important factor influencing the tree species richness in our study area. This is supported by Wulf and Naaf (2009), who reported a very similar observation from a managed forests. Fragmentation of natural forests due to anthropogenic pressure is a common phenomenon (Chazdon 2003). The present study area has been broken into patches of varying sizes separated by human disturbances such as fuel wood collection, bamboo collection, illicit felling and grazing (Mukul et al. 2014). Our result revealed that patch area had much pronounced effect on species richness both at tree, sapling and seedling level. In their studies Connell (1989), Dalling et al. (2002) found factors like surrounding vegetation, environmental modification, disturbance levels and dispersal limitation influence the species richness. Our findings also supported by the study of Singh (2002). Tripathi et al. (2010) in their study found a fairly large number of species with good regeneration performance in large forest fragments. However, the percentage of those species that showed fair, poor or no regeneration progressively increased with the increase of fragment size. Our study showed that species richness increased with disturbances in the forest. Interestingly, we found more species richness in sites with high level of disturbances. This finding was not supported by Bongers et al. (2009), who found disturbance contribute little to tree diversity. The major disturbances could also occur on a spatial scale that is larger or smaller than the 1 ha scale (Pickett and White 1985). Sagar et al. (2003) argued that if environmental change produced by disturbance is large, it may become lethal to greater numbers of established species than are, or can be, immediately replaced by immigrants. Furthermore, Collins et al. (1995) found a significant monotonic decline in species richness with increasing frequency of disturbance. According to Connell (1978), species richness is likely to be highest at the intermediate level of disturbance. This is not supported in our study. Our result also explored that tree species richness increases with tree cover in the forest. As our result showed more species richness with higher level of tree cover, and required both for sapling and seedling species to survive, it was strongly supported by the findings of Tripathi et al. (2010). This finding also supported the variation in the plant water availability within forest due to local differences in rainfall, dry season length, soil and/or topography (Bongers et al. 2009). Moreover, the richness of plant species due to their different responses to abiotic factors such as differential light levels, nutrient availability, water availability, wind and temperature. The abundance and richness of plants are also influenced by biotic factors such as birds, mammals and bats (Ramadhanil et al. 2008).

Result of constrained and unconstrained ordination revealed that tree species richness were explained by the disturbance, patch area and tree cover percentage. Similar findings was also reported by Pare et al. (2009).Yu and Sun (2013) in their study found that, the most important variable influencing species richness were strongly related to disturbance intensity and canopy openness. Sagar et al. (2003) found that habitat conditions and disturbance regime are important variable in controling the composition and richness of forest tree species in an area.

## 5 Conclusion

Our study indicated that level of forest disturbances, increased due to forest fragmentation in the area. The number of tree species was increased with increase in disturbance level. We also found that there was a strong correlation with forest patches, disturbance and tree cover and species richness. The patches, disturbance and tree cover contributed and favored the coexistence of particular tropical forest species. Such findings has an implication for the effective management or restoration of forest tree species with conservation significance, especially in highly disturbed and human modified landscapes. Forest patch leads to a reduction in total forest area, isolation of smaller patches in habitats and an increase in the disturbance level. All these can influence tree species richness in the smaller forest patch. Large patches were less disturbed than small forest patches and favored species which were not present in small patches. The small patches, however, favored regeneration of some species, which were absent in the large patches. Thus, both small and large patches together enhanced total tree species richness with patch area. Protective buffer of edge species around newly fragmented forest patches may useful to protect the species diversity in core forest and/or conservation zone. Moreover, we found both sapling and seedling can survive with high level of tree cover and disturbance. Future conservation efforts should address the broad ecological processes associated with other possible environmental correlates. We believe our study on the impact of selected site parameters on species richness will provide an important guideline for the future conservation planning not only in Bangladesh but also in other areas with similar bio-physical contexts.

## Acknowledgement

The comments from two anonymous reviewers and editor substantially improved the standard of our manuscript. Fieldwork for this study was supported by the Department of Forestry and Environment Science, Shahjalal University of Science and Technology, Bangladesh. Mohammad Belal Uddin was supported by a grant from University Research Center, Shahjalal University of Science and Technology to accomplish the field work for thi study. The first author would like to thanks Md. Abdullah-Al-Mamun and Syed Ajijur Rahman.

---

**Annex 1.** Tree species recorded from the study area

| Local name | Botanical name | Family | Abundance | Rank |
| --- | --- | --- | --- | --- |
| Agar | *Aquilaria malaccensis* Lamk. | Thymeleaceae | 4 | 79 |
| Akasmoni | *Acacia auriculiformis* A. Cunn. ex Benth | Leguminosae | 11 | 32 |
| Amloki | *Phyllanthus emblica* L. | Euphorbiaceae | 4 | 80 |
| Amm | *Mangifera indica* L. | Anacardiaceae | 8 | 47 |
| Aolagota | *Phyllanthus emblica* L. | Euphorbiaceae | 1 | 117 |
| Arjun | *Terminalia arjuna* (Roxb. ex DC.) Wight & Arn. | Combretaceae | 2 | 107 |
| Ata | *Annona muricata* L. | Annonaceae | 5 | 72 |
| Awal | *Vitex pubescens* Vahl. | Verbenaceae | 6 | 68 |
| Batna | *Quercus spicata* Smith | Fagaceae | 5 | 71 |
| Belam | *Averrho bilimbi* L. | Oxalidaceae | 6 | 63 |
| Belpoi | *Elaeocarpus floribundus* Bl. | Elaeocarpaceae | 8 | 48 |
| Bhadruk | *Vitex glabrata* R.Br. | Lamiaceae | 9 | 45 |
| Bogamedula | *Taphrosia candida* DC. | Fabaceae | 27 | 4 |
| Bohera | *Terminalia bellirica* (Gaertn.) Roxb. | Combretaceae | 22 | 11 |
| Bolas | *Sapium baccatum* | Euphorbiaceae | 6 | 61 |
| Bon badam | *Sterculia foetida* L. | Malvaceae | 1 | 113 |
| Bon chalta | *Leea crispa* L. | Leeaceae | 2 | 109 |
| Bon kadam | *Anthocephalus chinensis* (Lam.) A. Rich ex Walp. | Rubiaceae | 3 | 92 |
| Bon supari | *Areca triandra* Roxb. ex Buch.-Ham. | Arecaceae | 9 | 44 |
| Bonak | *Crataeva nurvosa* | Capparidaceae | 9 | 41 |
| Bonboroi | *Zizyphus oenoplea* (L.) Mill. | Rhamnaceae | 11 | 28 |
| Bor mala | *Calycarpa arborea* Roxb. | Verbenaceae | 2 | 98 |
| Bot | *Ficus benghalensis* L. | Moraceae | 6 | 57 |
| Buti dumur | *Ficus lamponga* Miq. | Moraceae | 16 | 17 |
| Chagalnadi | *Lepisanthes rubiginosa* (Roxb.) Leenh. | Sapindaceae | 5 | 73 |
| Chalta | *Dillenia indica* L. | Dilleniaceae | 4 | 88 |
| Champa | *Michelia champaca* L. | Magnoliaceae | 9 | 43 |
| Chapalish | *Artocarpus chaplasha* Roxb. | Moraceae | 35 | 1 |
| Chatim | *Alstonia scholaris* (L.) R. Br. | Apocynaceae | 14 | 19 |
| Chaulmoogra | *Hydnocarpus kurzii* (King) Warb. | Flacourtiaceae | 5 | 75 |
| Chengpisla | *Litsea glutinosa* (Lour.) Robinson | Lauraceae | 1 | 126 |
| Chikrasi | *Chickrassia tabularis* Wight & Arn. | Meliaceae | 11 | 27 |
| Chimta | Unknown |  | 4 | 87 |
| Deowa cham | *Artocarpus lacucha* Buch.-Ham. | Moraceae | 5 | 76 |
| Deua | *Artocarpus lacucha* Roxb. | Moraceae | 8 | 50 |
| Dhaki jam | *Syzygium firmum* Thw. | Myrtaceae | 4 | 83 |
| Dhup | *Canarium resiniferum* Bruce ex King | Burseraceae | 2 | 102 |
| Dumur | *Ficus hispida* L. f. | Moraceae | 7 | 51 |
| Ekus | *Dillenia scabrella* (D.Don) Roxb. ex Wall. | Dilleniaceae | 2 | 105 |
| Eucalyptus | *Eucalyptus camaldulensis* Dehnh. | Myrtaceae | 9 | 38 |
| Gab | *Diospyros malabarica* (Desr.) Kostel. | Ebenaceae | 3 | 93 |
| Gamar | *Gmelina arborea* Roxb. | Verbenaceae | 7 | 54 |
| Garjan | *Dipterocarpus turbinatus* Gaertn. | Dipterocarpaceae | 4 | 77 |
| Golap jam | *Syzygium jambos* (L.) Alston | Myrtaceae | 4 | 84 |
| Golsagu | *Caryota urens* L. | Arecaceae | 3 | 95 |
| Gora neem | *Melia azedarach* L. | Meliaceae | 7 | 56 |
| Goruch | Unknown |  | 6 | 67 |
| Gulli | Unknown |  | 26 | 6 |
| Guralebu | *Citrus limon* (L.) Osbeck | Rutaceae | 4 | 89 |
| Guti kadam | *Neonauclea sessilifolia* (Roxb.) Merr. | Rubiaceae | 2 | 104 |
| Haigakok | *Aegle marmelos* (L.) Corr. | Rutaceae | 12 | 25 |
| Hargoza | *Acanthus ilicifolius* L. | Acantheceae | 11 | 29 |
| Hargoza | *Dillenia pentagyna* Roxb. | Dilleniaceae | 4 | 85 |
| Holdu | *Adina cordifolia* (Roxb.) Hook. f. | Rubiaceae | 14 | 18 |
| Horitoki | *Terminalia chebula* Retz. | Combretaceae | 1 | 116 |
| Jalpoi | *Elaeocarpus robustus* Roxb. | Elaeocarpaceae | 3 | 96 |
| Jambura | *Citrus maxima* (Burm.) Merr. | Rutaceae | 2 | 99 |
| Janli badam | *Sterculia villosa* Roxb | Sterculiaceae | 4 | 78 |
| Jaowa | *Holigarna longifolia* Buch.-Ham. ex Roxb. | Anacardiaceae | 3 | 94 |
| Jarul | *Lagerstroemia speciosa* (L.) Pers. | Lythraceae | 11 | 26 |
| Jiga/ Jiol | *Lannea coromandelica* (Houtt.) Merr. | Anacardiaceae | 5 | 70 |
| Jog dumur | *Ficus racemosa* L. | Moraceae | 21 | 12 |
| Kadam | *Neolamarckia cadamba* (Roxb.) Bosser | Rubiaceae | 1 | 112 |
| Kaiphah | *Carica papaya* L. | Caricaceae | 4 | 86 |
| Kaju badam | *Anacardium occidentale* L. | Anacardiaceae | 13 | 21 |
| Kakra | *Glochidion multi-loculare* (Roxb. ex Willd) Muell.-Arg. | Euphorbiaceae | 22 | 8 |
| Kalo koro | *Albizia lebeck* (L.) Benth. | Leguminosae | 10 | 37 |
| Kalo jam | *Syzygium cacuminis* (Craib) Chantaran. & J.Parn. | Myrtaceae | 11 | 33 |
| Kanak | *Schima wallichii* Choisy | Theaceae | 7 | 53 |
| Kanthah | *Artocarpus heterophyllus* Lamk. | Moraceae | 8 | 49 |
| Kata kakra | *Lophopetalum fimbriatum* Wight | Celastraceae | 1 | 115 |
| Kata singra | *Castanopsis armata* (Roxb.) Spach | Fagaceae | 9 | 42 |
| Kath badam | *Terminalia catappa* L. | Combretaceae | 2 | 100 |
| Kau | *Garcinia cowa* Roxb. | Clusiaceae | 13 | 22 |
| Khami-1 | *Castanopsis tribuloides* (Sm.) A.DC. | Fagaceae | 20 | 13 |
| Khami-2 | *Lithocarpus elegans* Hook. f. | Fagaceae | 4 | 82 |
| Kharapat | *Garuga pinnata* Roxb. | Burseraceae | 18 | 15 |
| Khudi jam | *Syzygium fruticosum* (Roxb.) DC. | Myrtaceae | 22 | 10 |
| Koro | *Albizia chinensis* (Osbeck) Merr. | Leguminosae | 11 | 30 |
| Kumbi | *Carya arborea* Roxb. | Lecythidaceae | 3 | 91 |
| Kurchi | *Holarrhena antidysentica* (L.) Wall. | Apocynaceae | 5 | 74 |
| Lal dumur | *Erythrina resupinata* Roxb. | Leguminosae | 19 | 14 |
| Lal kakra | *Bruguiera gymnorhiza* | Rhizophoraceae | 12 | 24 |
| Lal menda | *Erythrina crista-galli* L. | Leguminosae | 2 | 101 |
| Lali | *Amoora wallichii* King | Meliaceae | 2 | 108 |
| Lohakat | *Xylia dolabriformis* Benth. | Leguminosae | 6 | 58 |
| Lalawal | *Vitex peduncularis* Wall. ex Schauer | Verbenaceae | 6 | 59 |
| Lotkon | *Baccaurea ramiflora* Lour. | Euphorbiaceae | 6 | 65 |
| Ludh | *Ilex godajam* Colebr. ex Hook.f. | Aquifoliaceae | 27 | 5 |
| Lu dijaylla | *Litsea lancifolia* (Roxb. ex Nees) Fern.-Vill. | Lauraceae | 1 | 128 |
| Mahogani | *Swietenia mahagoni* (L.) Jacq. | Meliaceae | 3 | 97 |
| Mailla | *Vallarix solanaceae* (Roth) Kuntze | Apocynaceae | 6 | 62 |
| Malakana koro | *Paraserianthes falcataria* (L.) I.C.Nielsen | Leguminosae | 13 | 20 |
| Mandar | *Erythrina variegata* L. | Leguminosae | 4 | 90 |
| Mander | *Erythrina orientalis* (L.) Merr. | Leguminosae | 1 | 124 |
| Mangium | *Acacia mangium* Willd. | Leguminosae | 1 | 120 |
| Menda | *Litsea monopetala* (Roxb.) Pers. | Lauraceae | 4 | 81 |
| Minjiri | *Cassia siamea* Lam. | Leguminosae | 10 | 35 |
| Moskon | *Pterospermum acerifolium* (L.) Willd. | Malvaceae | 25 | 7 |
| Nagersar | *Mesua nagesarium* (Burm. f.) Kosterm. | Clusiaceae | 1 | 121 |
| Neor | *Protium serratum* (Wall. ex Colebr.) Engl. | Burseraceae | 16 | 16 |
| Palm | *Elacis guineensis* Jacq. | Arecaceae | 10 | 36 |
| Panbot | *Ficus religiosa* L. | Moraceae | 2 | 110 |
| Pani jam | *Syzygium formosanum* (Hayata) Mor. | Myrtaceae | 2 | 111 |
| Para dumur | *Ficus auriculata* Lour. | Moraceae | 1 | 114 |
| Pataharolla | *Aporosa dioica* (Roxb.) Muell.-Arg | Euphorbiaceae | 1 | 118 |
| Ping | *Cynometra polyandra* Roxb. | Leguminosae | 2 | 103 |
| Pisli | *Aphanamixis polystachya* (Wall.) R.Parker | Meliaceae | 32 | 2 |
| Pitraj | *Aphanamixis polystachya* (Wall.) Parker | Meliaceae | 6 | 60 |
| Rokton | *Lophopetalum fimbriatum* Wight | Celastraceae | 1 | 130 |
| Rongi rata | *Aglaia spectabilis* (Miq.) S.S.Jain & S.Bennet | Meliaceae | 8 | 46 |
| Sada awal | *Vitex diversifolia* Kurz ex C.B. Clarke | Lamiaceae | 22 | 9 |
| Sada dumur | *Ficus hispida* L.f. | Moraceae | 11 | 31 |
| Sada hingra | *Cynometra ramiflora* | Caesalpiniaceae | 9 | 40 |
| Sal | *Shorea robusta* Gaertn. | Dipterocarpaceae | 6 | 66 |
| Shegun | *Tectona grandis* L.f. | Lamiaceae | 1 | 122 |
| Shaora | *Streblus asper* Lour. | Moraceae | 9 | 39 |
| Shet Shimul | *Ceiba pentandra* | Bombacaceae | 7 | 55 |
| Silkoro | *Albizia procera* (Roxb.) Benth. | Leguminosae | 11 | 34 |
| Simul | *Bombex ceiba* L. | Malvaceae | 7 | 52 |
| Siris | *Albizia lebbeck* (L.) Benth. | Leguminosae | 31 | 3 |
| Supari | *Areca catechu* L. | Arecaceae | 2 | 106 |
| Tejpata | *Cinnamomum tamala* (Buch.-Ham.) T.Nees & Eberm. | Lauraceae | 1 | 129 |
| Tel gorjan | *Dipterocarpus turbinatus* C.F. Gaertn. | Dipterocarpaceae | 1 | 119 |
| Telsur | *Hopea odorata* Roxb. | Dipterocarpaceae | 1 | 127 |
| Tentul | *Tamarindus indica* L. | Caesalpiniaceae | 1 | 125 |
| Thona | *Oroxylum indicum* (L.) Kurz | Bignoniaceae | 5 | 69 |
| Tipali | *Trewia nudiflora* L. | Euphorbiaceae | 1 | 123 |
| Toon | *Toona ciliata* M. Roem. | Meliaceae | 6 | 64 |
| Udal | *Sterculia colorata* Roxb. | Sterculiaceae | 12 | 23 |

**Annex 2.**
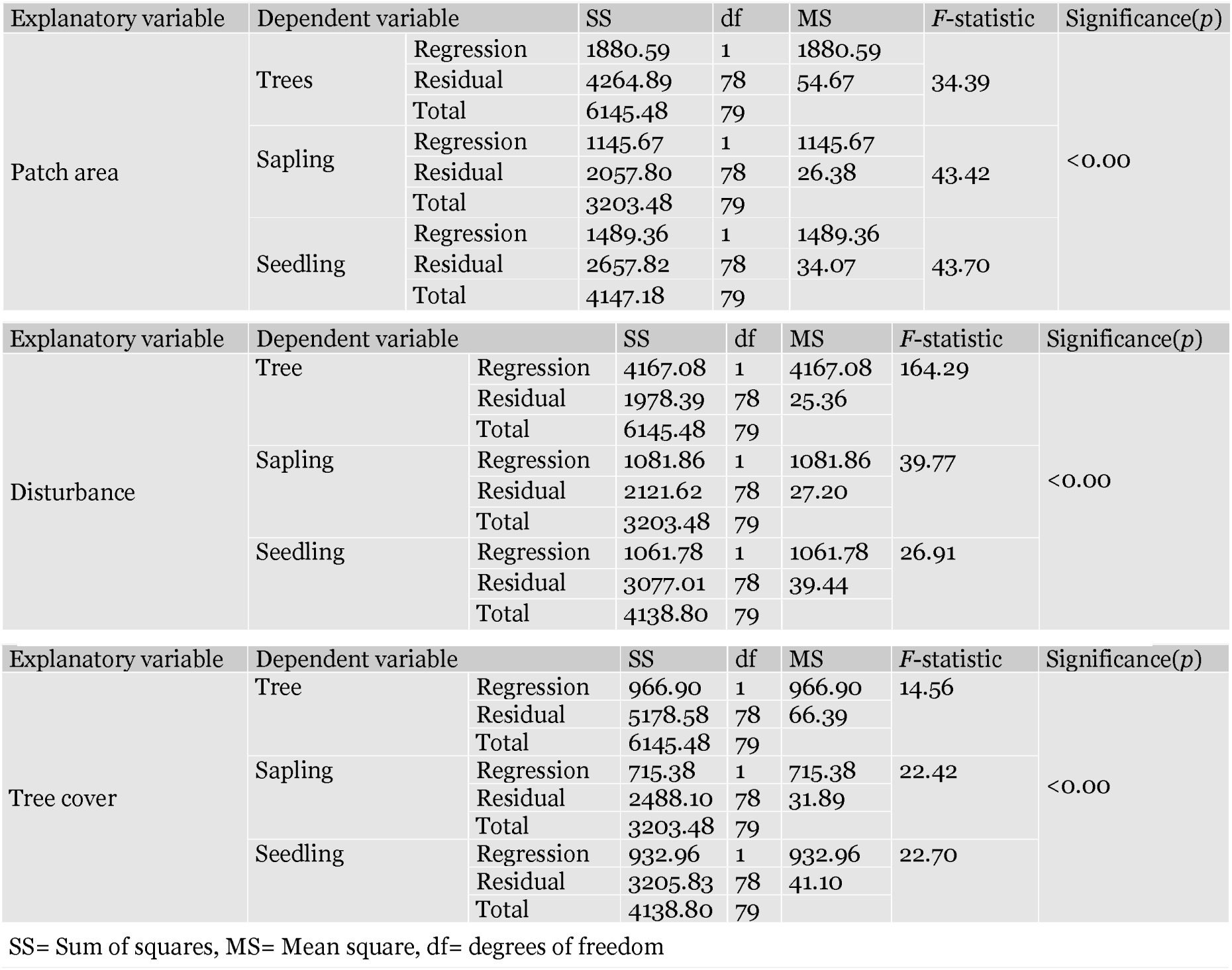
Analysis of variance (ANOVA) summary between dependent and independent variable. **Table A** Analysis of variance (ANOVA) of linear regression of patch area on species richness **Table B** Analysis of variance (ANOVA) of linear regression model between disturbance and species richness **Table C** Analysis of variance <u>(ANOVA)</u> of linear regression model between tree cover and species richness

**Supplementary material 1.**
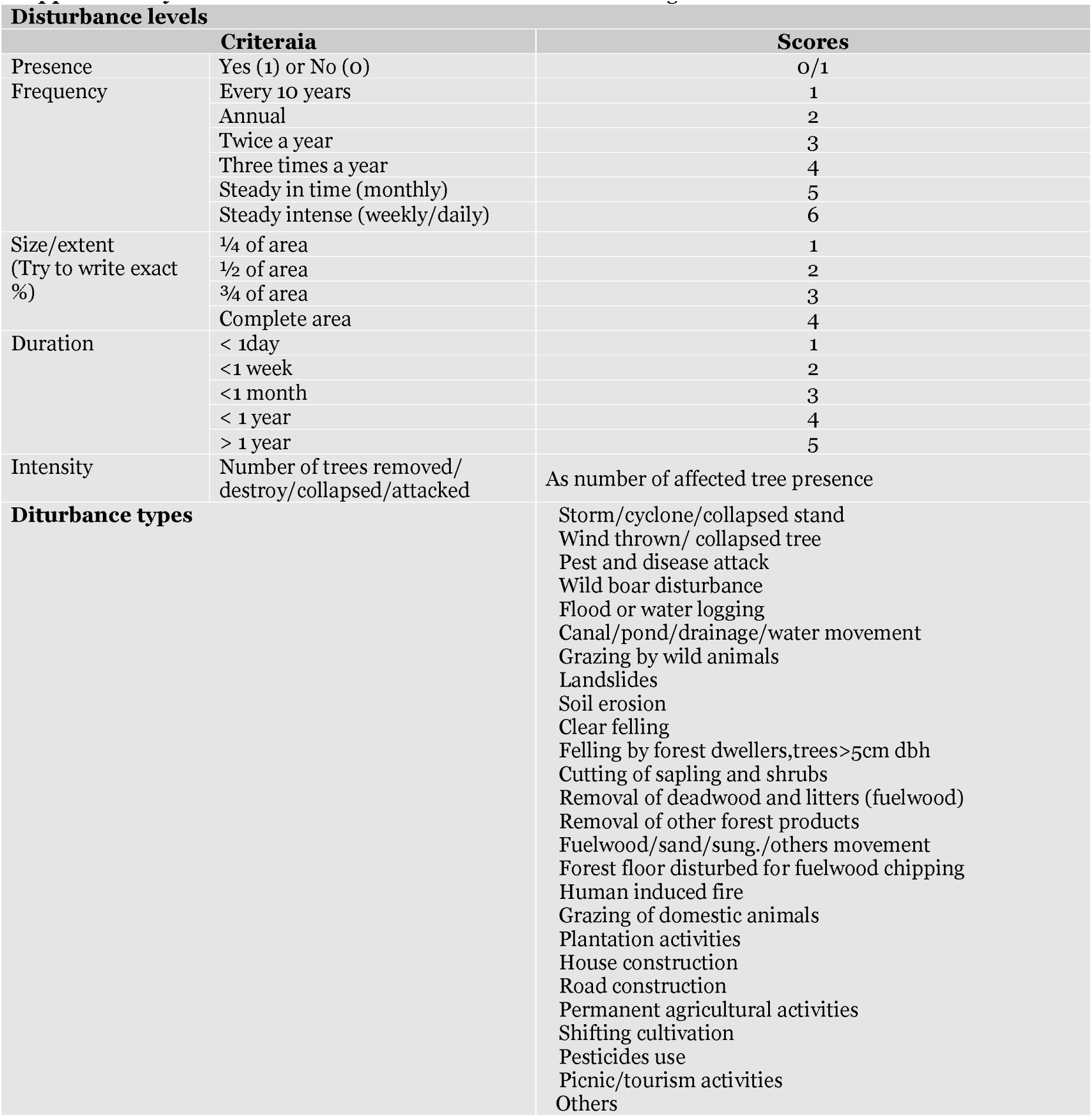
Checklist of the disturbances and ranking criteria

## References

Acharya B (1999) Forest biodiversity assessment. a spatial analysis of tree species diversity in Nepal. ITC publication number 72. ITC, Enschede, The Netherlands.

Alamgir M, Mukul SA, Turton S (2015) Modelling spatial distribution of critically endangered Asian elephant and Hoolock gibbon in Bangladesh forest ecosystems under a changing climate. Applied Geography 60: 10-19. DOI: 10.1016/j.apgeog.2015.03.001

Biohabitats T (2012) Forest ecological assessment. Virginia Tech. Baltimore, Maryland.

Bongers F, Poorter L, Hawthorne WD, Sheil D (2009) The intermediate disturbance hypothesis applies to tropical forests, but disturbance contributes little to tree diversity. Ecology Letters 12: 798-805. DOI: 10.1111/j.1461-0248.2009.01329.x

Brokaw N (1998) Fragments past, present and future. Tree 13: 382-383. DOI: 10.1111/j.14610248.2009.01329.x

Burslem DFRP, Whitmore TC (1999) Species diversity, susceptibility to disturbance and tree population dynamics in tropical rain forest. Journal of Vegetation Science 10: 767-776. DOI: 10.2307/3237301

Chazdon RL (2003) Tropical forest recovery: legacies of human impact and natural disturbances. Perspectives in Plant Ecology, Evolution and Sytematics 6: 51-71. DOI: 10.1078/1433-8319-00042

Collins SL, Glenn SM, Gibson DJ (1995) Experimental analysis of intermediate disturbance and initial floristic composition: decoupling cause and effect. Ecology 76: 486-492. DOI: org/10.2307/1941207

Connell JH (1978) Diversity in tropical rain forests and coral reefs. Science 24: 1302-1310. DOI: 10.1126/science.199.4335.1302

Connell JH (1989) Some processes affecting the species composition in forest gaps. Ecology 70: 560-562. DOI. org/10.2307/1940205

Chowdhury MSH, Koile M (2010) An overview on the protected area system for forest conservation in Bangladesh. Journal of Forest Research 21:111-118.

Dalling JW, Muller-Landau HC, Wright SJ, Hubbell SP (2002) Role of dispersal in the recruitment limitation of neotropical pioneer species. Journal of Ecology 90:714-727.

Ehrlich PR, Wilson EO (1991) Biodiversity studies: science and policy. Science 253: 758-762. DOI: 10.1126/science.253.5021.758

Forman RTT (1995) Land mosaics, the ecology of landscapes and regions. Cambridge University Press. Cambridge, UK. p632.

Galanes IT, Thomlinson JR (2008) Relationships between spatial configuration of tropical forest patches and woody plant diversity in northeastern Puerto Rico. Plant Ecology 201:101-113. DOI: 10.1007/978-90-481-2795-5_9

GOB (2010) Forest department official website. Government of People’s Republic of Bangladesh, Dhaka, Bangladesh. http://www.bforest.gov.bd/land.php, accessed on February 15, 2015

Holdgate M (1996) The ecological significance of biological diversity. Ambio 25: 409-416.

Khan MASA, Uddin MB, Uddin MS, Chowdhury MSH, Mukul SA (2007) Distribution and status of forests in the tropics: Bangladesh perspective. Proc. Pakistan Academy Science 44:145-153.

Kibria MG, Rahman SA, Imtiaj A, Sunderland T (2011) Extent and consequences of tropical forest degradation: successive policy options for Bangladesh. Journal of Agricultural Science and Technology 1:29-37.

Laurance WF, Bierregaard JRO (1997) Tropical forest remnants: ecology, management and conservation of fragmented communities. University of Chicago Press. Chicago and London.

Mukul SA (2008) The role of traditional forest practices in enhanced conservation and improved livelihoods of indigenous communities: case study from Lawachara National Park, Bangladesh. 1^st^ International Conference on ‘Forest Related Traditional Knowledge and Culture in Asia. Seoul, Korea. pp24-28.

Mukul SA (2014) Biodiversity conservation and ecosystem functions of traditional agroforestry systems: case study from three tribal communities in and around Lawachara National Park. In: Chowdhury MSH (ed.), Forest Conservation in Protected Areas of Bangladesh - Policy and Community Development Perspectives. Springer Switzerland. pp171-179.

Mukul SA, Herbohn J (2016) The impacts of shifting cultivation on secondary forests dynamics in tropics: a synthesis of the key findings and spatio temporal distribution of research. Environmental Science & Policy 55: 167-177. DOI: 10.1016/j.envsci.2015.10.005

Mukul SA, Herbohn J, Rashid AZMM, Uddin MB (2014) Comapring the effectiveness of forest law enforcement and economic incentive to prevent illegal logging in Bangladesh. International Forest Review 16: 363-375. DOI: 10.1505/146554814812572485

Mukul SA, Rashid AZMM, Quazi SA, et al. (2012) Local peoples’ response to co-management regime in protected areas: a case study from Satchari National Park, Bangladesh. Forests, Tress and Livelihoods 21: 16-29. DOI: 10.1080/14728028.2012.669132

Mukul SA, Rashid AZMM, Uddin MB (2012) The role of spiritual beliefs in conserving wildlife species in religious shrines of Bangladesh. Biodiversity 13: 108-114. DOI: 10.1080/14888386.2012.694596

Murcia C (1995) Edge effects in fragmented forests: implications for conservation. Trends in Ecology & Evolution 10:58-62. DOI: 10.1016/S0169-5347(00)88977-6

NSP (2006) Management plans for Lawachara National Park. Nishorgo Support Project (NSP), Bangladesh.

Pandey HN, Tripathi OP, Tripathi RS (2003) Ecological analysis of forest vegetation of Meghalaya. In: Bhatt et al. (eds.), Approaches for increasing agricultural Productivity in Hill and Mountain ecosystem. ICAR Barapani, Meghalaya, India.

Pare S, Savadogo P, Tigabu M, et al. (2009) Regeneration and spatial distribution of seedling populations in Sudanian dry forests in relation to conservation status and human pressure. Tropical Ecology 50: 339-353.

Pickett STA, White PS (eds.) (1985) The ecology of natural disturbance and patch dynamics. Academic Press. San Diego.

Raghubanshi AS, Tripathi A (2009) Effect of disturbance, habitat fragmentation and alien invasive plants on floral diversity in dry tropical forests of Vindhyan highland: a review. Tropical Ecology 50: 57-69.

Ramadhanil R, Tjitrosoedirdjo SS, Setiadi D (2008) Structure and composition of understory plant assemblages of six land use types in the Lore Lindu National Park, Central Sulawesi, Indonesia. Bangladesh. Association of Plant Taxonomists 15: 1-12. DOI: 10.3329/bjpt.v15i1.911

Rao P, Barik SK, Pandey HN, et al. (1990) Community composition and tree population structure in a sub-tropical broad-leaved forest along a disturbance gradient. Vegetation 88:151-162. DOI: 10.1007/BF00044832

Rahman SA, Baldauf C, Mollee EM, et al. (2013) Cultivated plants in the diversified homegardens of local communities in Ganges valley, Bangladesh. Science Journal of Agricultural Research & Management 2013:1-6. DOI: 10.7237/sjarm/197

Rahman SA, Foli S, Pavel MAA, et al. (2015) Forest, trees and agroforestry: better livelihoods and ecosystem services from multifunctional landscapes, International Journal of Development and Sustainability 4: 479-491.

Sagar R, Raghubanshi AS, Singh JS (2003) Tree species composition, dispersion and diversity along a disturbance gradient in a dry tropical forest region of India. Forest Ecology and Management 186: 61-71. DOI: 10.1016/S0378-1127(03)00235-4

Shrestha TK (1999) Nepal country report on biological diversity. Katmandu, IUCN, Nepal. p133.

Singh JS (2002) The biodiversity crisis: a multifaceted review. Current Science 82: 638-647.

Sohel MSI, Mukul SA, Burkhard B (2015) Landscape’s capacities to supply ecosystem services in Bangladesh: a mapping assessment for Lawachara National Park. Ecosystem Services 12: 128-135. DOI: 10.1016/j.ecoser.2014.11.015

Tilman D (2000) Causes, consequences and ethics of biodiversity. Nature 405: 208-211. DOI: 10.1038/35012217

Tripathi OP, Upadhaya K, Tripathi RS, Pandey HN (2010) Diversity, dominance and population structure of tree species along fragment-size gradient of a subtropical humid forest of northeast India. Environmental and Earth Sciences 2: 97-105.

Uddin MB, Steinbauer MJ, Jentsch A, et al. (2013) Do environmental attributes, disturbances and protection regime determine the distribution of exotic plant species in Bangladesh forest ecosystems? Forest Ecology and Management 303: 72-80. DOI: 10.1016/j.foreco.2013.03.052

Vieira DLM, Scariot A (2006) Principles of natural regeneration of tropical dry forests for restoration. Restoration Ecology 14:11-20. DOI: 10.1111/j.1526-100X.2006.00100.x

Whittaker RH (1972) Evolution and measurement of species diversity. Taxon 21: 213-251. DOI: 10.2307/1218190

Wulf M, Naaf T (2009) Herb layer response to broadleaf tree species with different leaf litter quality and canopy structure in temperate forests. Journal of Vegetation Science 20: 517-526. DOI: 10.1111/j.1654-1103.2009.05713.x

Yu M, Sun OJ (2013) Effects of forest patch type and site on herb-layer vegetation in a temperate forest ecosystem. Forest Ecology and Management 300:14-20. DOI: 10.1016/j.foreco.2012.12.039

